# Regulation of tumor proliferation by unlocking silent synapses on metastatic cancer cells

**DOI:** 10.64898/2026.02.16.706220

**Authors:** Aiden J. Houcek, Ihsan Alp Uzay, Lisa M. Monteggia, Amanda Linkous, Barbara Fingleton, Ege T. Kavalali

**Affiliations:** Department of Pharmacology, Vanderbilt University, Nashville, TN, USA; Vanderbilt Brain Institute, Vanderbilt University, Nashville, TN, USA; Department of Biological Sciences, Vanderbilt University, Nashville, TN, USA

**Author notes:** These authors contributed equally (Aiden J. Houcek, Ihsan Alp Uzay). Corresponding author: Ege T. Kavalali.

## Abstract

Several studies have revealed deleterious synapse formation onto cancer cells within the brain tumor microenvironment, yet these synapses are ∼100-fold weaker in presynaptic release rates and postsynaptic strength relative to *bona fide* synapses formed between neurons. Here, we find that most of the functional synapses on tumor cells are kept dormant and can be unlocked by overcoming GABA_B_ receptor-mediated metabotropic signaling in neurons. Scavenging G_βγ_ signaling in neurons increased presynaptic release probability on tumor cells and augmented cancer cell proliferation. Optical analysis of the tumor microenvironment revealed regulated secretion of neurotransmitters from tumor cells in response to GABA_B_ receptor inhibition or electrical stimulation. These results reveal how cancer cells with a high propensity for brain metastasis leverage precise moments of aberrant excitation between neurons to engage reciprocal interactions that ultimately fuel cancer proliferation.

## Introduction

Certain cancers share a uniquely high propensity for brain metastasis. Small cell lung cancer, melanoma, and breast cancer are among the most likely peripheral tumors to metastasize to the brain in humans^1,2^. While distinct in tissues of origin and non-neoplastic counterparts, common symptoms emerge following brain metastasis in humans^3,4^. Neuronal hyperexcitability and altered synaptic transmission have been documented as common phenotypes in the tumor microenvironment across various primary and metastatic cancers in the brain^5–10^. While indirect interactions such as physical compression and edema in surrounding brain tissue undoubtedly contributes to symptoms experienced by patients with brain tumors, functional integration through direct synapse formation on tumor cells suggests that dysregulated neuronal communication in the tumor microenvironment may be a result of precise interactions that engage specific and actionable circuits and signaling mechanisms^11–14^. Brain metastatic cancers also promote dysregulated neuronal activity via secretion of various signaling molecules such as synaptogenic factors, as well as neurotransmitters such as glutamate, GABA, and others^15–17^. In line with this evidence of the secretory properties of various cancers, recent studies reveal an intrinsic excitability or action potential-like waveforms in both small cell lung cancer and glioma^18–20^. These findings suggest that both primary brain tumors and cancers with a high propensity for brain metastasis may engage secretory properties in addition to the currently established postsynaptic integration of these cancer types.

While recent evidence has uncovered both cell autonomous and circuit-dependent signaling mechanisms that contribute to the integration of cancer cells with neuronal networks^21,22^, the direct functional consequences of elevated neuronal activity on cancer cell pathophysiology remains largely unexplored. Here, we set out to understand the effects of rapidly augmenting neuronal activity on tumor cells. We found that inducing aberrant excitation between neurons and tumor cells activated a group of dormant synapses on cancer cells that are typically suppressed at basal levels of neuronal activity. Comparing neuron-neuron synaptic activity in the same conditions revealed that cancer cells receive a filtered and downscaled version of the same synaptic activity pattern between neurons. Visualizing tumor cell vesicle trafficking revealed a secretory phenotype of brain adapted and neuroendocrine cancers when grown with neurons. Secreted molecules specifically from breast and lung cancer cells generated currents on neurons which further promoted aberrant excitation. Together, these results reveal a rapid and reciprocal positive feedback loop that promotes tumor cell proliferation in moments of dysregulated neuronal activity and suggest potential therapeutic strategies to alleviate seizure-induced cancer growth in brain metastasis.

## Results

### Unlocking spontaneous synaptic currents on tumor cells

We began by exploring how augmenting neuronal activity affects tumor cell physiology by adding human neuroendocrine small cell lung cancer (H69, SCLCA, referred to as H69), BRAF V600E melanoma (SK-MEL-5, referred to as SK-5), or triple negative brain breast cancer cells (MDA-231-BrM2, referred to as BrM2) to hippocampal neuronal networks^23,24^ (Fig. 1A). We allowed tumor cells to integrate with neurons for at least 10 days prior to conducting experiments. To induce neuronal hyperactivity in this system, we perfused the GABA_A_ receptor antagonist bicuculline and recorded synaptic activity from neurons or tumor cells in the network using whole-cell patch clamp electrophysiology. Isolated inhibitory postsynaptic currents (IPSCs) confirmed that bicuculline inhibits GABAergic transmission and subsequently disinhibits glutamatergic transmission between neurons (Fig. S1A and B). We then recorded spontaneous currents from tumor cells before and after bicuculline perfusion. Here, we detected minimal spontaneous currents at baseline in cancer cells, consistent with previous studies employing patch-clamp recordings of glioma cells and small cell lung cancer cells^5,7,12^. Interestingly, bicuculline application induced a robust increase in spontaneous currents in H69 SCLC, SK-MEL-5, and BrM2 cancer cells (Fig. 1B and 1C). The frequency of these bicuculline-induced spontaneous currents in tumors was approximately 1 Hz and presented in a burst-like pattern of activity. These currents were silenced by the AMPA receptor antagonist CNQX as well as the voltage-gated sodium channel blocker tetrodotoxin (TTX) (Fig. S1C). This rapid induction of synaptic transmission on tumor cells led us to hypothesize that presynaptic inhibition of neurotransmission on tumor cells is mediated by a form of tonic metabotropic signaling (Fig. 1D). We explored this hypothesis by pharmacologically inhibiting GABA_B_ receptors with the competitive antagonist saclofen while recording spontaneous activity on cancer cells^25^. Remarkably, saclofen application produced an identical current pattern in tumor cells to bicuculline perfusion (Fig. 1E, F and S1D). These currents were also sensitive to CNQX and were not augmented further following bicuculline application (Fig. 1G), suggesting that bicuculline and saclofen converge on their mechanism of action. Like bicuculline, saclofen perfusion also augmented spontaneous synaptic transmission between neurons (Fig. S1E). Comparing neuron-neuron synaptic activity to neuron-tumor activity following this unsilencing revealed that tumor cells receive a filtered and downscaled version of the same synaptic activity patterns between neurons (Fig. S2A and B). Taken together, these findings suggest that augmenting spontaneous neuronal activity unlocks silent synapses on tumor cells by overcoming G-protein coupled metabotropic inhibition of neurotransmitter release.

**Fig. 1.**
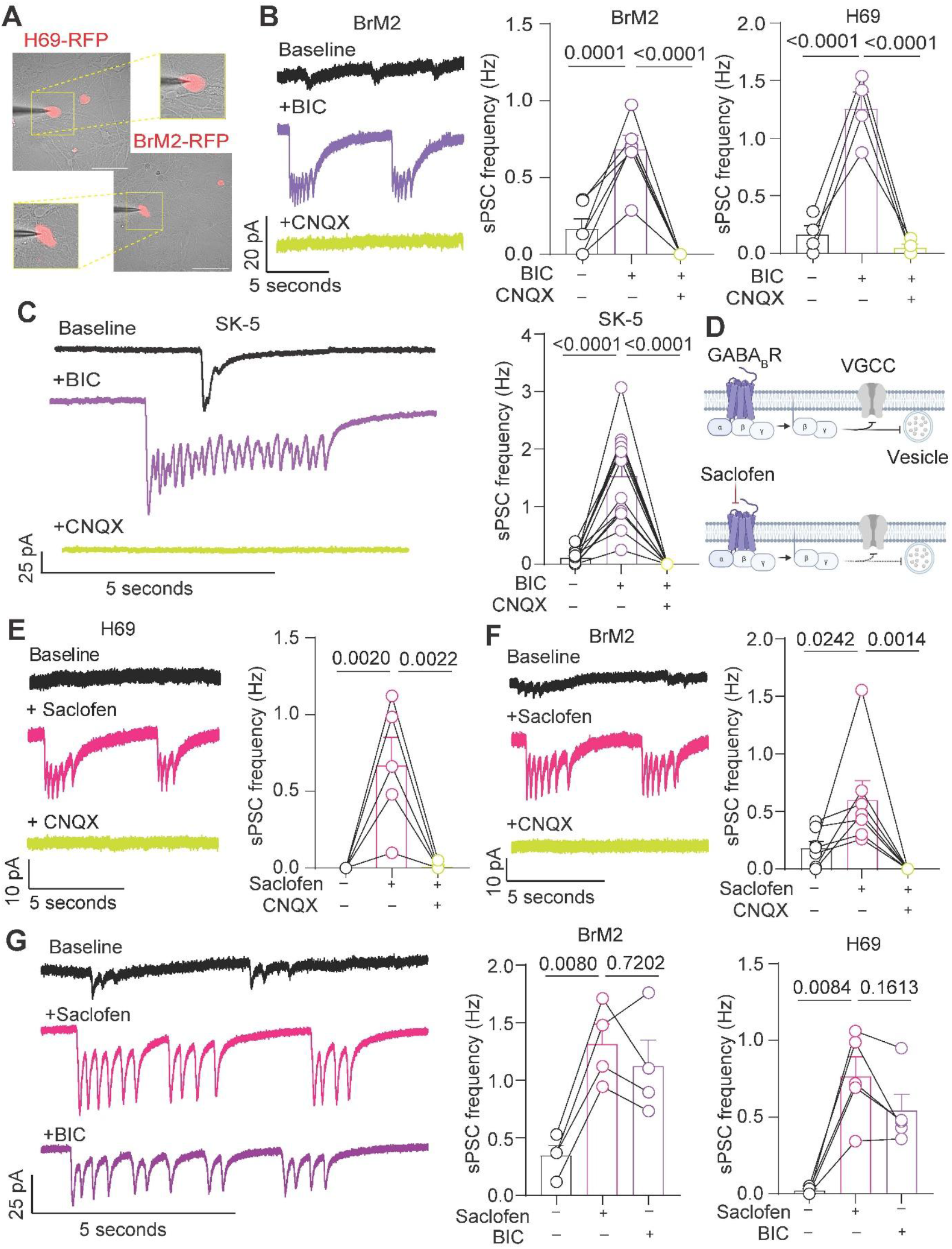
Blocking GABA_A_ or GABA_B_ receptors unlocks spontaneous synaptic currents in tumor cells co-cultured with primary hippocampal neurons. **A**. Fluorescent and DIC images with magnified insets of patched H69-RFP and BrM2-RFP tumor cells in co-culture with rat hippocampal neurons. Scale bar = 100µm. **B**. Representative trace and quantification of spontaneous synaptic current frequency in H69 and BrM2 cells following bicuculline and CNQX application in co-culture with neurons. **C**. Example traces and quantification of sPSC frequency after bicuculline and CNQX perfusion in SK-5 cells co-cultured with hippocampal neurons. **D**. Illustration of GABA_B_ receptor-mediated inhibition of presynaptic neurotransmission. **E** and **F**. Representative traces and quantification of spontaneous synaptic currents in H69 (E) and BrM2 (F) cells co-cultured with neurons following saclofen and CNQX perfusion. **G**. Representative trace and quantification of spontaneous synaptic currents in H69 and BrM2 cells co-cultured with neurons following saclofen and bicuculline application. Statistical significance was assessed using one-way ANOVA with Tukey’s multiple comparisons (B,C,E,F,G).

### Presynaptic inhibition of tumoral neurotransmission by GABA_B_ receptors

The rapid activation of spontaneous neurotransmission on cancer cells following GABA_A_ or GABA_B_ receptor inhibition suggests that a subclass of functional synapses are formed on tumor cells but inhibited under resting conditions of synaptic transmission between neurons. This hypothesis led us to test if synaptic transmission is suppressed around tumor cells. To answer this question, we recorded spontaneous presynaptic calcium transients in neurons using GCaMP8s tethered to the presynaptic SNARE protein Synaptobrevin2 (Syb2-GcAMP8s). This probe allowed us to specifically visualize synaptic calcium transients in neuronal synapses both proximal to tumor cells as well as distal regions where no tumor cells were present in the same system (Fig. 2A). Imaging spontaneous presynaptic calcium transients in the proximity of cancer cells revealed a reduction in both calcium transient frequency and amplitude relative to more distal presynaptic terminals (Fig. 2B and S3A). As expected, saclofen application produced a robust increase in the frequency of presynaptic calcium transients at single synapses both distal and proximal to tumor cells (Fig. 2C), consistent with our functional analysis. To uncover the relative levels of neurotransmitter secretion following GABA_B_R inhibition at presynaptic terminals in proximity to tumor cells, we expressed a Syb2-Super ecliptic pHluorin (Syb2-SEP) in neurons and measured SEP signal intensity at neuronal synapses distal and proximal to tumor cells. Here, we found an increase in peak SEP signal intensity at presynaptic terminals in proximity to cancer cells following saclofen application (Fig. 2D and S3B), suggesting that the chronic suppression of synaptic activity around cancer cells elicits a larger reversal in synaptic disinhibition during moments of aberrant excitation. Ammonium chloride, which leads to alkalinization of secretory vesicles and subsequently induces SEP fluorescence, further revealed multiple synaptic contacts on tumor cells (Fig. S3E). Given this evidence for GABA_B_R-mediated inhibition of neurotransmission around cancer cells, we next employed a molecular genetic approach to impair GABA_B_ receptor signaling in neurons by expressing a myristoylated c-terminal fragment of GPCR kinase 2 (myr-ct-GRK2) to scavenge G_βγ_ signaling^26^ (Fig. 2E and F). Here, we found that scavenging G_βγ_ signaling in neurons robustly augmented the amplitude of postsynaptic currents on tumor cells and increased presynaptic release probability (Fig. 2G and H). Repetitive stimulation at 20Hz in G_βγ_ scavenger conditions also revealed a marked increase in short-term synaptic depression on tumor cells (Fig. 2I). These functional consequences of scavenging G_βγ_ signaling in neurons were consistent across SK-5 melanoma and neuron-neuron synapses (Fig. S4A-F). Furthermore, we found that scavenging G_βγ_ signaling elevated spontaneous neurotransmission at baseline (Fig. 3A and B) and occluded the induction of spontaneous transmission on tumor cells following saclofen perfusion (Fig. 3C, D and S3C and D). We found that these functional phenotypes in response to scavenging G_βγ_ signaling or saclofen incubation correlated with an increase in tumor cell proliferation (Fig. 3E and F). Taken together, these results demonstrate that tonic GABA_B_ receptor signaling actively suppresses synaptic transmission on tumor cells and inhibits cancer cell proliferation at baseline levels of neurotransmission.

**Fig. 2.**
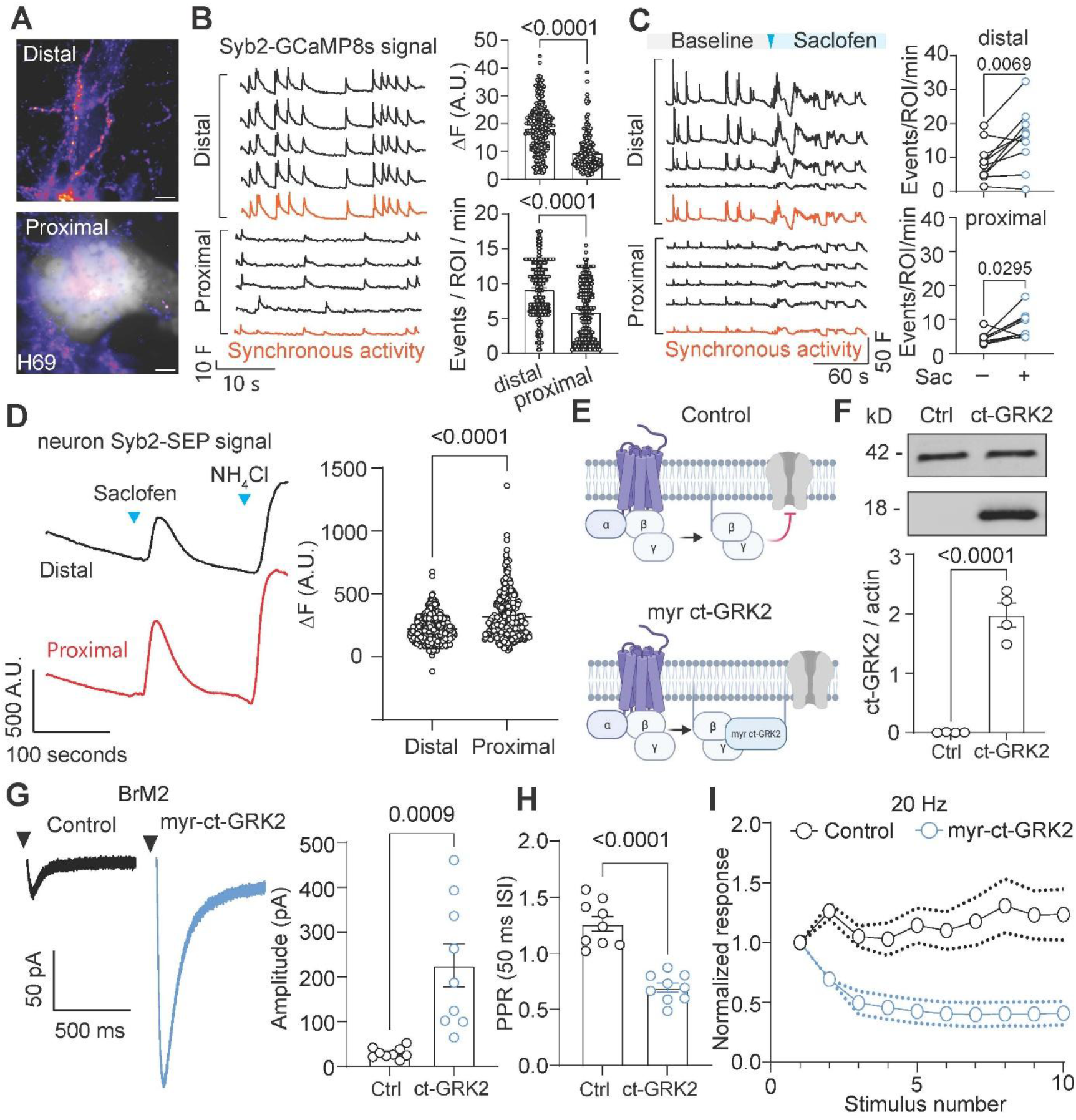
Presynaptic terminals are chronically suppressed around tumor cells and can be unsilenced by scavenging G_βγ_ signaling in neurons. **A**. Example image of Syb2-GCaMP8s expression in neurons distal and proximal to H69 tumor cells in co-culture with neurons. Scale bar = 5 µm. **B**. Representative recordings and quantification of spontaneous Syb2GCaMP8s signal frequency and amplitude from individual ROIs at presynaptic terminals distal and proximal to BrM2 cancer cells. **C**. Example recording and quantification of Syb2-GCaMP8s signal frequency in presynaptic terminals distal and proximal to tumor cells before and after saclofen application. **D**. Example trace and quantification of Syb2-SEP signal expressed in neurons at presynaptic terminals distal or proximal to H69 tumor cells. **E**. Schematic illustrating the mechanism of myr-ct-GRK2 mediated scavenging of G_βγ_ signaling. **F**. Western blot and quantification of ct-GRK2 expression relative to β-actin in hippocampal neuron cultures. **G**. Representative trace and quantification of ePSC amplitude in BrM2 cells co-cultured with neurons transduced with myr-ct-GRK2 relative to control. **H**. Quantification of paired pulse ratio using a 50 ms interstimulus interval. **I**. Normalized ePSC amplitudes from control and myr-ct-GRK2 conditions as a function of stimulus number at 20Hz stimulation frequency. Statistical significance was assessed using paired samples t-test (C), independent samples t-test (D,F,G,H).

**Fig. 3.**
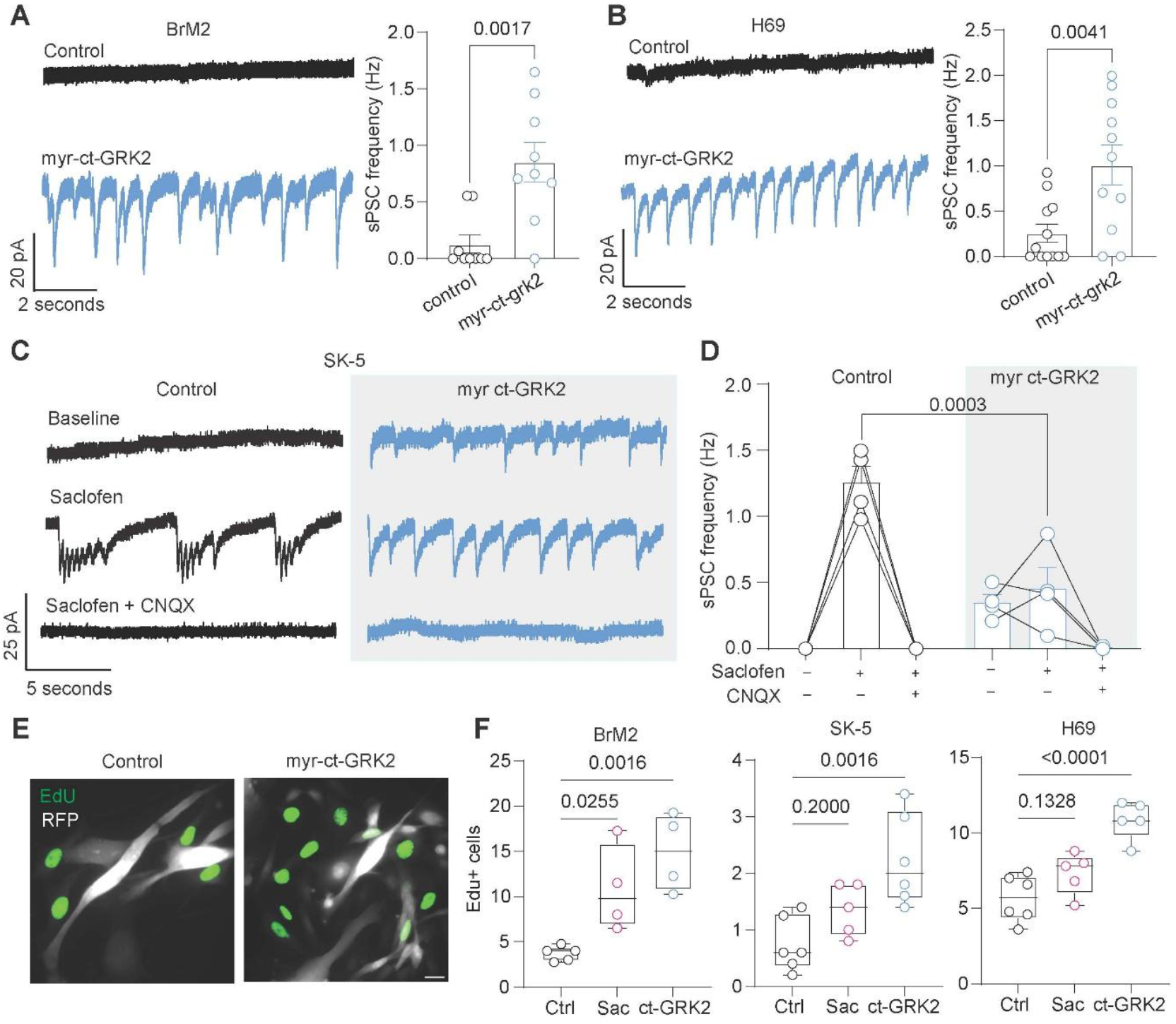
Scavenging G_βγ_ signaling in neurons occludes saclofen-induced spontaneous neurotransmission and increases tumor cell proliferation. **A** and **B**. Example traces and quantification sPSC frequency in BrM2 and H69 SCLC cells with hippocampal neurons expressing myr-ct-GRK2 relative to control. **C** and **D**. Example traces and quantification of sPSC amplitude in SK-5 cells co-cultured with neurons expressing myr-ct-GRK2 following saclofen and CNQX perfusion. **E**. Example image of EdU+ tumor cells in coculture with hippocampal neurons expressing myr-ct-GRK2 relative to control conditions. Scale bar = 20 µm. **F**. Quantification of EdU+ tumor cells in response to 24-hour saclofen incubation or myr-ct-GRK2 expression in neurons. Statistical significance was assessed using independent samples t-test (A,B,D), One-way ANOVA with Dunnett’s multiple comparisons (F).

### Regulated secretion from brain adapted tumor cells

Cancer cells and non-neoplastic counterparts are known to secrete various molecules to regulate proliferation, paracrine signaling, and migration^27–29^. Recent research studying glioma as well as small cell lung cancer have revealed action potential-like waveforms in these tumor cells in response to depolarization, potentially indicating a form of secretory function in response to sufficient excitation^5,18–20^. However, whether this depolarization in tumor cells is coupled to regulated secretion, as in neurons, remains unknown. To uncover if presynaptic properties of tumor cells are coupled to neuronal activity, we expressed Syb2-SEP in cancer cells to visualize secretion during moments of augmented synaptic activity between neurons (Fig. 4A). Remarkably, high frequency electrical stimulation of neuronal networks revealed a robust secretory phenotype from tumor cells that was not observed in cancer cell monocultures (Fig. 4B). This form of regulated secretion was observed in approximately 55% of H69 and BrM2 cells cultured with neurons, whereas SK-5 cells rarely exhibited a secretory phenotype in response to stimulation of surrounding neurons (Fig. 4C and D). As a positive control, we expressed Syb2-SEP in neurons as previously described which revealed robust and reliable activity in response to high frequency stimulation (Fig. 4E). Saclofen application also resulted in a robust increase in Syb2-SEP signal in H69 and BrM2 cells co-cultured with neurons (Fig. 4F). We then asked if this secretory phenomenon of brain adapted tumor cells persists in a more translational model of brain metastasis. To test this, we performed intracardiac injections of BrM2 cells expressing Syb2-SEP in mice. We confirmed brain metastasis in these mice using IVIS imaging and prepared acute brain slices to record secretory properties of metastatic cells following saclofen application (Fig. 4G). Similar to our *in vitro* experiments, saclofen perfusion resulted in a significant increase in BrM2 Syb2-SEP fluorescence in *ex vivo* brain slices (Fig. 4H and I). Ammonium chloride also augmented SEP intensity in metastatic BrM2 cells. Since saclofen does not readily permeate the blood brain barrier, we then employed the brain penetrant GABA_B_ receptor antagonist CGP 35348 to uncover the role of GABA_B_ receptor inhibition on BrM2 proliferation *in vivo*^30^. Functionally, we found that CGP generated similar phenotypes as saclofen on BrM2 cells cultured with hippocampal neurons (Fig. S4A and B). Acute perfusion of CGP also increased Syb2-SEP intensity in metastatic brain tumors (Fig. 4J) and elevated the proliferation of brain metastatic breast cancer cells *in vivo* (Fig. 4K). Taken together, these findings reveal how brain metastatic cancer cells gain secretory abilities in neuronal contexts to leverage moments of aberrant excitation for regulated exocytosis.

**Fig. 4.**
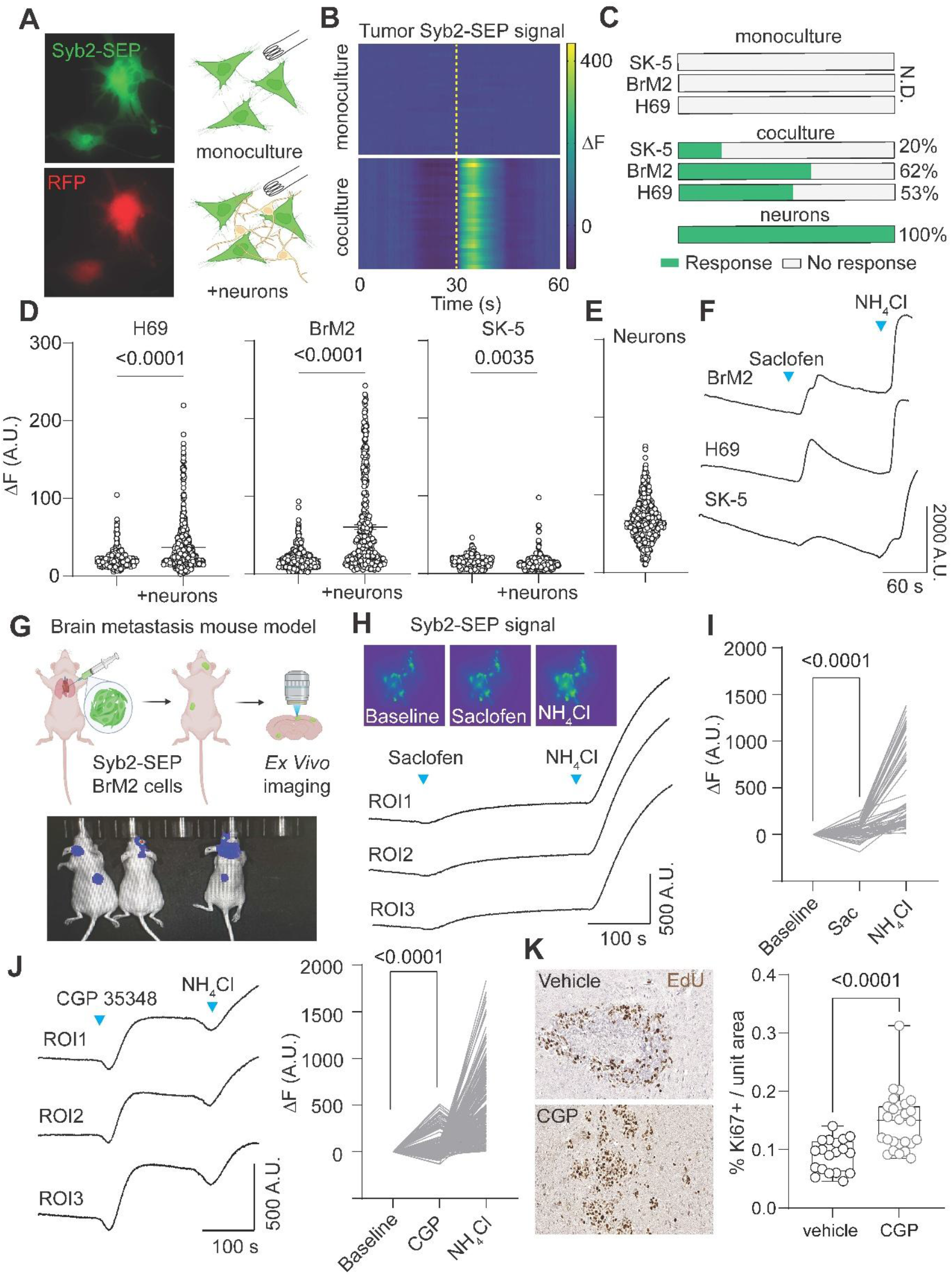
Electrical stimulation and inhibition of GABA_B_ receptors induces regulated secretion from tumor cells. **A**. Example fluorescent image of Syb2-SEP expression and RFP expression in tumor cells and schematic illustrating experimental strategy to test stimulation-dependent secretion in tumor cells co-cultured with neurons. Scale bar = 20 µm. **B**. Heat map showing tumor cell Syb2-SEP intensity in response to stimulation at 30 seconds in monoculture or co-cultured with neurons. Time of stimulation indicated by dashed yellow line. **C**. Percentage of responding tumor cells to electrical stimulation in monoculture or coculture relative to neurons expressing Syb2-SEP. **D**. Quantification of max Syb2-SEP intensity in tumor cells in monoculture or co-cultured with neurons in response to stimulation. **E**. Quantification of Syb2-SEP intensity in neurons in response to stimulation. **F**. Example recordings of Syb2-SEP intensity in BrM2, H69, and SK5 cancers cells following saclofen and ammonium chloride perfusion. **G**. Schematic illustrating brain metastasis mouse model using MDA-231-BrM2 tumor cells expressing Syb2-SEP and example image of confirmed brain metastasis in mice using IVIS. **H**. Example images and recording trace of BrM2 Syb2-SEP signal in BrM2 brain metastatic tumor cells. **I**. Quantification of Syb2-SEP signal intensity following saclofen or ammonium chloride perfusion. **J**. Example recordings and quantification of Syb2-SEP signal in brain metastatic BrM2 cells following CGP 35348 and ammonium chloride perfusion. **K**. Example images of brain metastatic breast cancer from nude mice treated with vehicle or CGP 35348 and quantification of Ki67+ tumor cells. Individual data points represent brain slices from 3-4 mice per condition. Statistical significance was assessed using independent samples t-test (D, K), Paired samples t-test (I,J).

### Reciprocal regulation of neuronal activity from tumor cells

Thus far, our results suggest that inhibiting GABA_B_ receptor signaling between neurons and tumor cells unlocks spontaneous synaptic transmission on cancer cells and simultaneously induces regulated exocytosis from H69 and BrM2 cancer cells. Therefore, we next asked what molecules are secreted from these tumor cells during moments of aberrant excitation and how these secreted factors may affect neuronal activity. To uncover what molecules are secreted by tumor cells, we incubated cancer cells with modified Tyrode’s containing calcium and magnesium for 12 hours. The conditioned Tyrode was then collected and filtered to remove cellular debris (Fig. 5A). Tumor cell conditioned Tyrode was then perfused on neurons in the presence of TTX to inhibit action potential-dependent network activity. As a negative control, unconditioned Tyrode solution was perfused first which did not generate detectable charge transfer in neurons. Perfusing BrM2 or H69 conditioned Tyrode on neurons generated a robust charge in voltage clamp configuration (Fig. 5B), whereas application of SK-5 conditioned Tyrode’s did not generate currents in neurons (Fig. 5C). We speculated that neurotransmitter secretion from H69 and BrM2 cells is partially responsible for the observed currents in neurons, as previous studies have documented secretory properties in glioma cells. When we pharmacologically isolated glutamatergic transmission in neurons, we once again observed robust charge transfer following BrM2 or H69 conditioned Tyrode perfusion (Fig. S6A and B), indicating that glutamate is indeed secreted by H69 and BrM2 cells. We also detected evidence of GABA secretion from H69 and BrM2 cells (Fig. S6C and D), supporting our previous findings of synaptic suppression at presynaptic terminals in proximity to tumor cells. Next, we acutely incubated cancer cells with modified Tyrode’s containing saclofen to address if the previously observed increase in Syb2-SEP intensity correlates with more secreted factors into the extracellular space. Here, we found that saclofen conditioned Tyrode from H69 and BrM2 cells resulted in significantly larger currents when perfused on neurons relative to acutely conditioned control Tyrode solution (Fig. 5D and E). Saclofen conditioned Tyrode from SK-5 cells did not generate currents in neurons (Fig. 5F), consistent with the markedly lower levels of secretion observed using Syb2-SEP as previously described. Importantly, saclofen was added to the control Tyrode in these experiments after harvesting to prevent confounding effects of saclofen perfusion on synaptic transmission. Next, to investigate how secreted molecules from tumor cells regulates neuronal activity, we conducted current clamp recordings of neurons and perfused tumor cell conditioned Tyrode while measuring spontaneous action potential firing. Here, we found that BrM2 and H69 conditioned Tyrode’s increased actional potential frequency in neurons and augmented the neuronal resting membrane potential (Fig. 5G and H). Consistent with the lack of charge transfer following SK-5 conditioned Tyrode perfusion, we did not observe a significant change in neuronal AP firing following SK-5 conditioned Tyrode perfusion (Fig. 5I). Together, these results show how secreted molecules from H69 and BrM2 tumor cells influence neuronal signaling and further promote aberrant excitation in moments of augmented synaptic transmission.

**Fig. 5.**
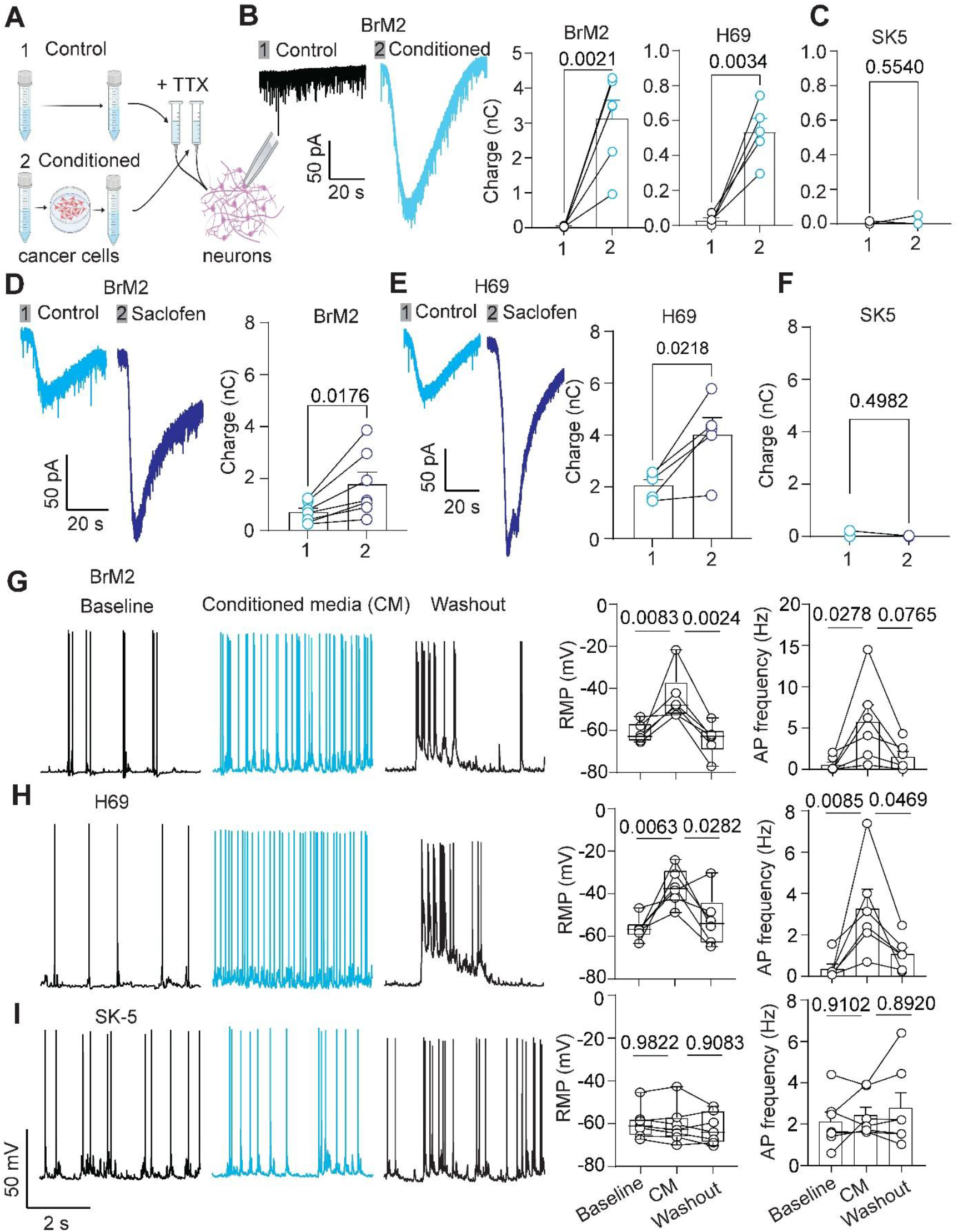
Acute perfusion of secreted factors from cancer cells generate glutamate and GABAergic currents and augment action potential firing in neurons. **A**. Schematic illustrating experimental strategy to test the effects of tumor cell secreted factors on neurons. **B**. Example trace and quantification of unconditioned and conditioned Tyrode perfusion from BrM2 and H69 cells on neurons in voltage clamp configuration. **C**. Quantification of charge in neurons following tumor cell conditioned Tyrode perfusion from SK-5 cells. **D** and **E**. Example traces and quantification of charge transfer in neurons following acutely conditioned Tyrode perfusion or saclofen conditioned Tyrode from BrM2 cells (D) or H69 cells (E) in neurons. **F**. Quantification of charge transfer in neurons following acutely conditioned Tyrode perfusion or saclofen conditioned Tyrode from SK-5 cells. **G-I**. Example traces and quantification of resting membrane potential and spontaneous action potential frequency in neurons following tumor cell conditioned media perfusion. Statistical significance was assessed using paired sampled t-test (B, C, D, E, F), One-way ANOVA with Tukey’s multiple comparisons (G, H, I).

## Discussion

The functional integration of cancer cells with neurons has been documented in multiple tumor types including glioma, SCLC, melanoma, and breast cancer^5,8,31,32^. Tracing studies have also revealed diverse neuronal inputs on tumor cells and a range of secreted neurotransmitters on xenografted brain tumors^33^. These studies and others have revealed evidence for GABAergic synapse formation on both diffuse midline glioma and SCLC cells^5,8,34,35^. Here, we reveal that blocking GABA_A_ receptors can also reveal a population of synapses on tumor cells that are not readily detected in single stimulation paradigms to study neuron-tumor synapses. Additionally, the difference in acute or chronic pharmacological manipulations may induce substantial changes in neuronal signaling and tumor cell proliferation, highlighting the importance of considering precise therapeutic windows in patients with brain tumors. Regardless, the chronic suppression of functional synapses on tumor cells and reciprocal secretion from brain adapted cancers described here reveals an additional layer to the therapeutic framework for targeting neuron-tumor electrochemical interactions. Future studies may elucidate how these moments of neuronal hyperactivity are either hijacked or generated by cancer cells to promote invasion in brain metastasis. In addition to the induction of spontaneous synaptic transmission on tumor cells following GABA_B_ receptor inhibition, we also describe how scavenging G_βγ_ signaling augments the amplitude of synaptic currents and induced a robust increase in release probability and short-term synaptic depression on tumor cells^36,37^. These findings highlight how a genetic manipulation can be employed to investigate the pathophysiology of synaptic plasticity on tumor cells and further underscore the role of GABA_B_R function in the vicinity of tumor cells.^38–40^. We then uncovered a regulated secretion from H69 and BrM2 tumor cells. While we describe neuron-dependent secretory properties in response to electrical stimulation of surrounding neurons, our data also illustrate a role of GABA_B_ receptor activity in inducing secretory phenotypes and suppressing baseline levels of tumor cell proliferation. While largely known to inhibit neurotransmission, GABA_B_ receptor function can also promote release in specific regions such as the medial habenula^41^. Previous studies have also described a positive role of GABA_B_ receptor activity on tumor cell proliferation through β-catenin signaling^42^, highlighting both circuit and microenvironment-dependent roles of GABA_B_ receptor signaling in brain tumor growth.

Our investigation of BrM2, H69, and SK5 tumor cells also revealed some significant distinctions in the presynaptic properties following integration with neurons. While all three cell lines functionally integrated with neurons, breast and lung cancer cells shared a secretory phenotype that was largely absent in SK5 melanoma cells. Previous studies have shown evidence for glutamate secretion from B16F1 melanoma cells^43^, suggesting heterogeneity in secretory properties within individual tumor types. Furthermore, secreted glutamate is often correlated with invasiveness in brain tumors^44^ and, although SK5 was originally isolated from a metastatic site (axillary lymph node), SK5 cells are considered less invasive compared to other melanoma lines such as A375 and A2058^45^. SK5 cells also exhibit reduced protein expression of annexin A1^45^ – a key player in the stabilization of enzymes that enhance glutamate production in some cancer models^46^. Thus, the inclusion of additional tumor cell models with varied levels of invasiveness and genotypic complexity will be important for future evaluation of secretory phenotypes. Together, our findings reveal how the vast majority of synapses on cancer cells are chronically suppressed and how cancer cells with a high propensity for brain metastasis leverage precise moments of aberrant excitation between neurons to rapidly engage reciprocal interactions that ultimately fuel cancer cell proliferation. These findings provide a methodological framework to study synaptic coupling of cancer cells to neurons and potential therapeutic strategies to prevent seizure-induced tumor growth in brain metastasis.

## Acknowledgements

We thank members of the Kavalali and Monteggia labs for their feedback on the manuscript. This work was supported by National Institute of Health grants NS134128 to ETK, MH070727 to LMM, and an Exceptional Project Award to B.F. from the Breast Cancer Alliance.

