## Supplementary Figures for "Regulation of tumor proliferation by unlocking silent synapses on metastatic cancer cells"

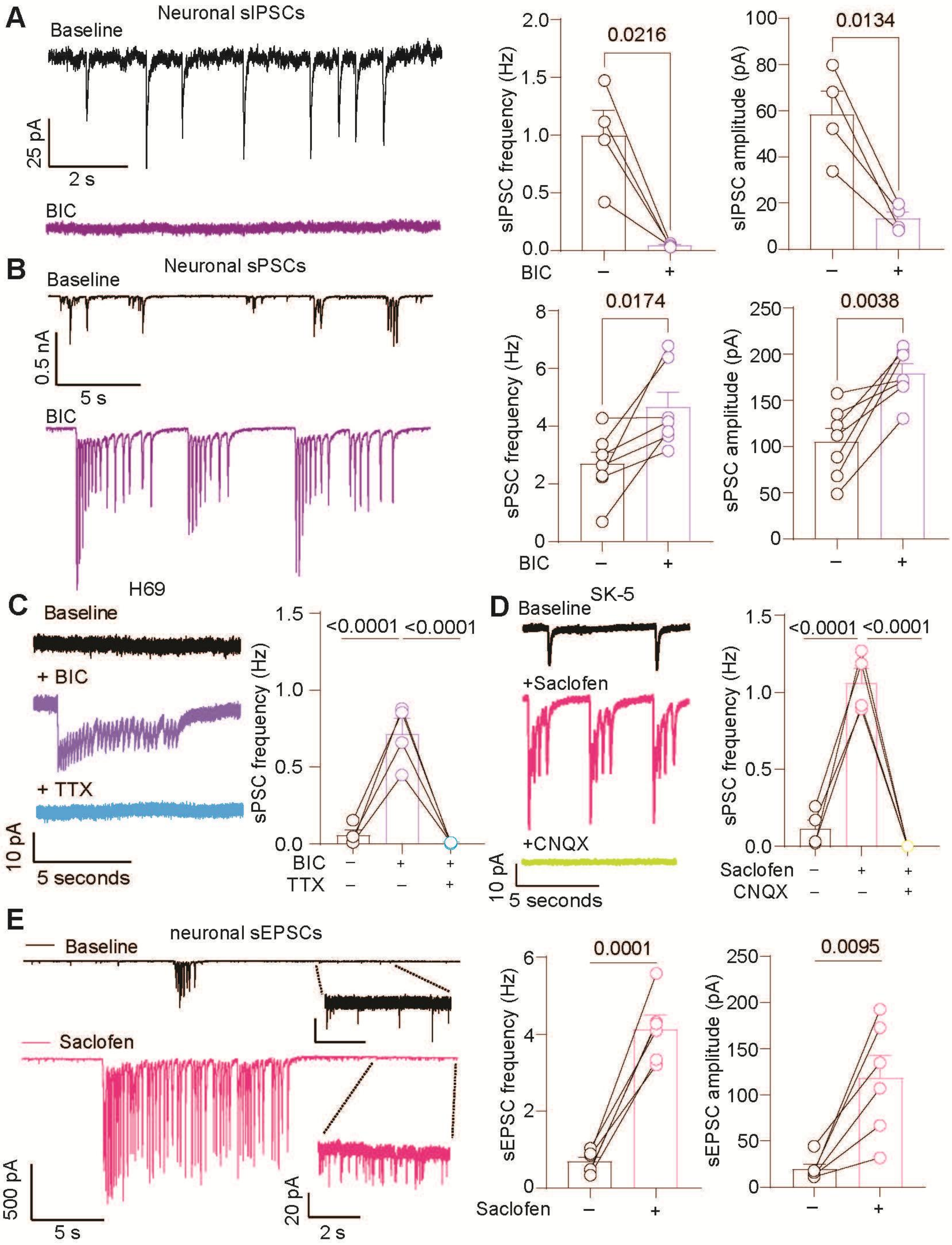

A
Neuronal sIPSCs
Baseline
25 pA
2 s
BIC
0.0216
sIPSC frequency (Hz)
0.0134
sIPSC amplitude (pA)
B
Neuronal sPSCs
Baseline
0.5 nA
5 s
BIC
0.0174
sPSC frequency (Hz)
0.0038
sPSC amplitude (pA)
H69
C
Baseline
+ BIC
+ TTX
10 pA
5 seconds
<0.0001
<0.0001
sPSC frequency (Hz)
BIC
TTX
SK-5
D
Baseline
+Saclofen
+CNQX
10 pA
5 seconds
<0.0001
<0.0001
sPSC frequency (Hz)
Saclofen
CNQX
E
neuronal sEPSCs
Baseline
Saclofen
500 pA
5 s
20 pA
2 s
0.0001
sEPSC frequency (Hz)
0.0095
sEPSC amplitude (pA)

**Fig. S1. Inhibiting $GABA_A$ or $GABA_B$ receptors induces aberrant synaptic transmission between neurons and cancer cells**. **A.** Example traces and quantification of neuronal sIPSC frequency and amplitude before and after bicuculline perfusion. **B.** Example traces and quantification of neuronal sPSC frequency and amplitude before and after bicuculline perfusion. **C.** Representative trace and quantification of spontaneous synaptic currents in H69 cells co-cultured with neurons after bicuculline and TTX application. **D.** Example traces and quantification of sPSC frequency in SK-5 cells after saclofen and CNQX application. **E.** Example recordings and quantification of sPSC frequency and amplitude in neurons following saclofen perfusion. Statistical significance was assessed using paired samples t-test (A,B,E), One-way ANOVA with Tukey's multiple comparisons (G, D).

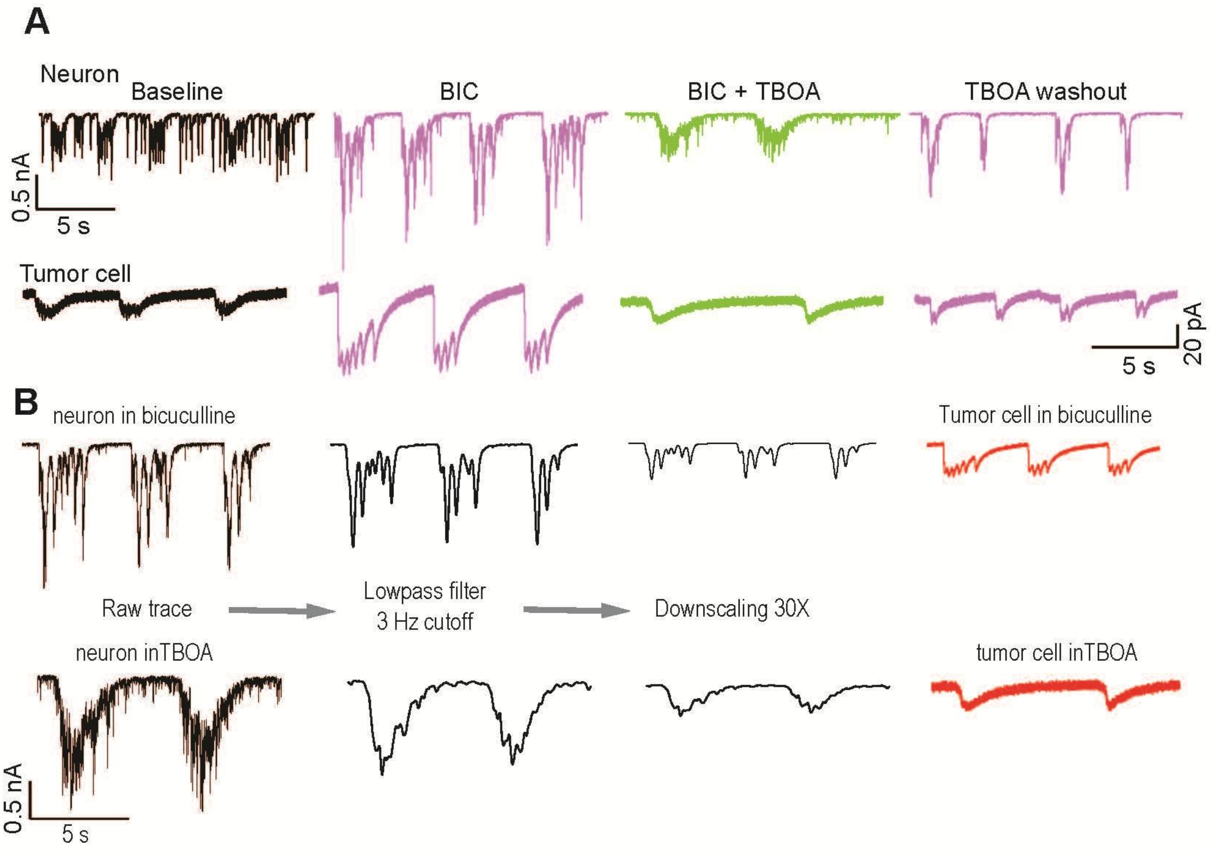
A
Neuron
Baseline
BIC
BIC + TBOA
TBOA washout
0.5 nA
5 s
Tumor cell
20 pA
5 s
B
neuron in bicuculline
Tumor cell in bicuculline
Raw trace
Lowpass filter
3 Hz cutoff
Downscaling 30X
neuron inTBOA
tumor cell inTBOA
0.5 nA
5 s

**Fig. S2. Tumor cells receive a filtered and downscaled version of synchronous synaptic transmission between neurons. A.** Example traces of neuronal synaptic transmission following bicuculline, TBOA, and TBOA + bicuculline washout (top traces). Bottom traces show synaptic currents recorded from an H69 tumor cell in the same pharmacological context as neurons. **B.** Example recordings of filtered and downscaled synaptic activity between neurons in bicuculline (top) or bicuculline + TBOA (bottom) relative to an example recording of synaptic events on an H69 tumor cell in the same conditions.

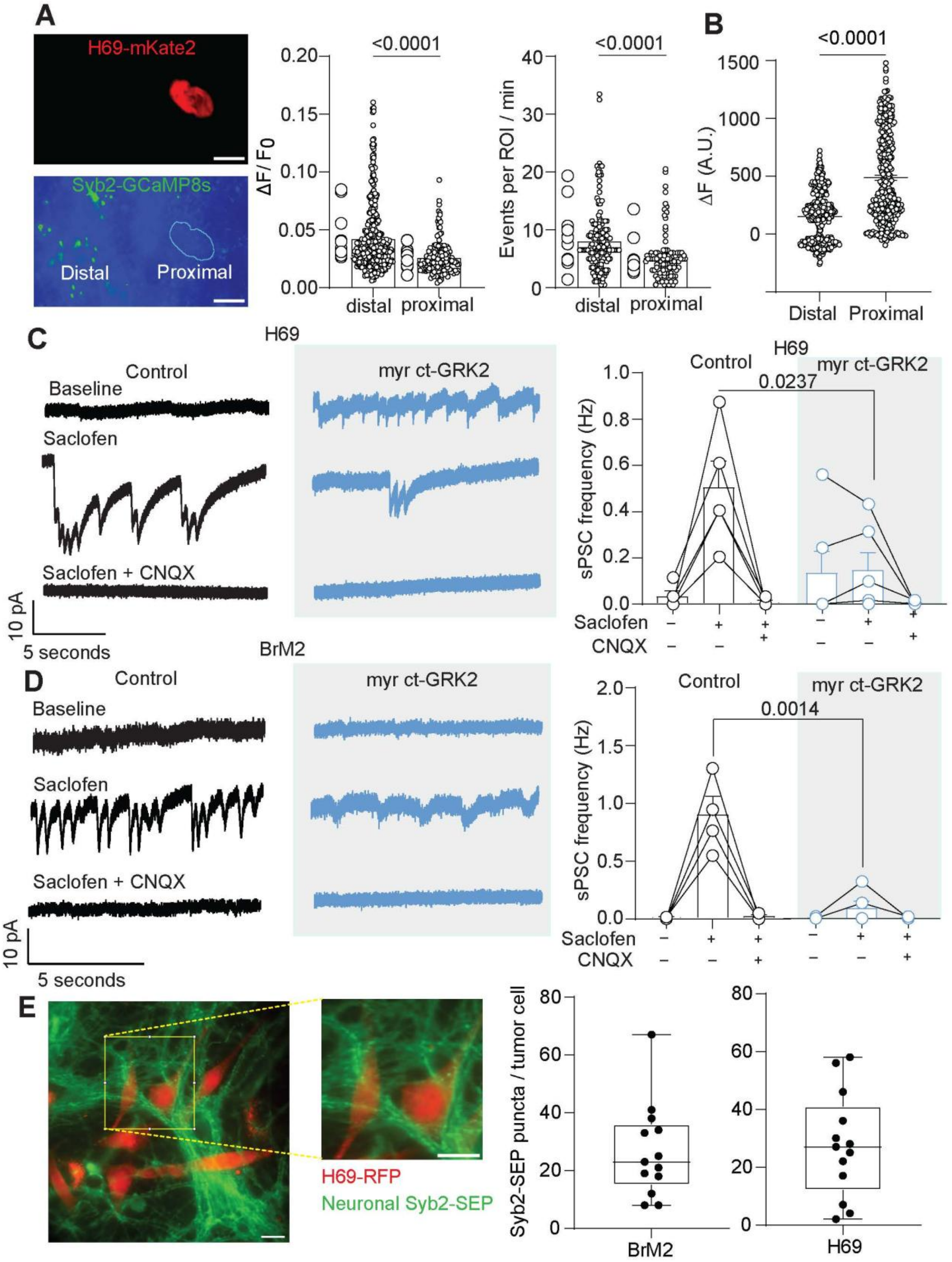
A
H69-mKate2
Syb2-GCaMP8s
Distal
Proximal
<0.0001
ΔF/F0
distal
proximal
Events per ROI / min
B
ΔF (A.U.)
Distal
Proximal
C
H69
Control
Baseline
Saclofen
Saclofen + CNQX
myr ct-GRK2
10 pA
5 seconds
sPSC frequency (Hz)
0.0237
CNQX
BrM2
D
0.0014
E
H69-RFP
Neuronal Syb2-SEP
Syb2-SEP puncta / tumor cell
BrM2
H69

**Fig. S3. Presynaptic suppression of neurotransmission around tumor inhibits cancer is occluded by scavenging $G_{\beta\gamma}$ signaling in neurons. A.** Example image and quantification of Syb2-GCaMP8s signal frequency an amplitude in neuronal synapses distal and proximal to H69 cancer cells. **B.** Quantification of peak neuronal Syb2-SEP signal following saclofen perfusion at synapses distal or proximal to BrM2 cells. **C** and **D.** Representative traces and quantification of sPSC frequency in H69 cells (C) or BrM2 cells (D) co-cultured with neurons transduced with myr-ct-GRK2 following saclofen and CNQX application relative to control conditions in the same pharmacological conditions. **E**. Fluorescent image and magnified inset of Syb2-SEP signal in neurons (green) following ammonium chloride application and H69 RFP (red) expressing tumor cells and quantification of Syb2-SEP puncta number on H69 and BrM2 tumor cells. Statistical significance was assessed using independent samples t-test (A, B, C, D).

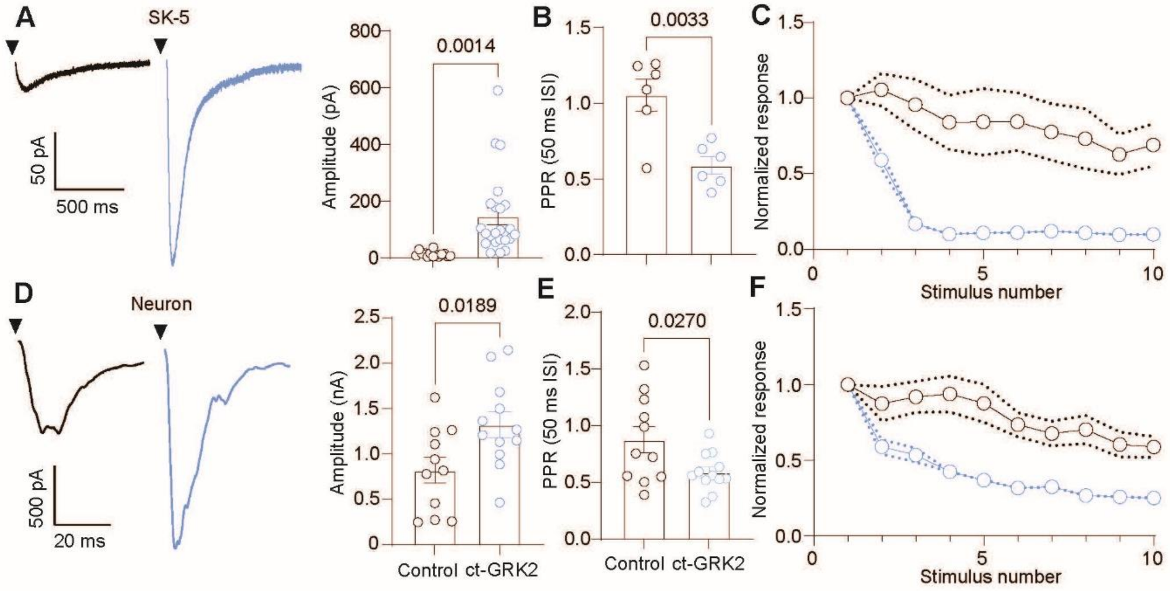

A
SK-5
50 pA
500 ms
Amplitude (pA)
0.0014
B
PPR (50 ms ISI)
0.0033
C
Normalized response
Stimulus number
D
Neuron
500 pA
20 ms
Amplitude (nA)
0.0189
E
PPR (50 ms ISI)
0.0270
F
Normalized response
Stimulus number
Control
ct-GRK2

**Fig. S4. Scavenging $G_{\beta\gamma}$ signaling increases presynaptic release probability in neurons and tumor cells. A** Representative trace and quantification of ePSC amplitude in SK-5 cells co-cultured with neurons transduced with myr-ct-GRK2 relative to control. **B**. Quantification of paired pulse ratio using a 50 ms interstimulus interval. **C.** Normalized ePSC amplitudes from control and myr-ct-GRK2 conditions as a function of stimulus number at 20Hz stimulation frequency. **D**. Representative trace and quantification of eEPSC amplitude in neurons expressing myr-ct-GRK2 relative to control. **E** and **F**. Quantification of paired pulse ratio and normalized amplitude as a function of stimulus number in neurons expressing myr-ct-GRK2 relative to control. Statistical significance was assessed using independent samples t-test (A, B, D, E).

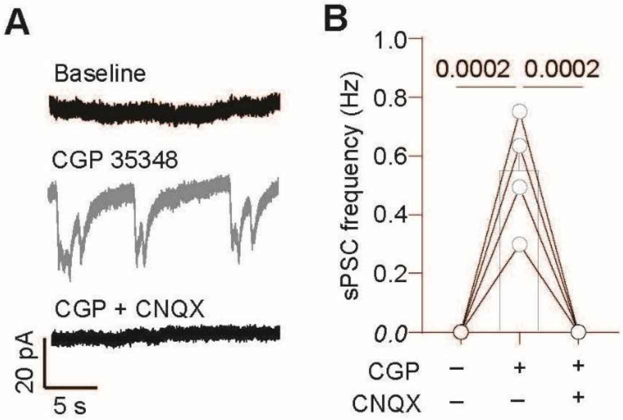

A
Baseline
CGP 35348
CGP + CNQX
20 pA
5 s
B
1.0
0.8
0.6
0.4
0.2
0.0
sPSC frequency (Hz)
0.0002
0.0002
CGP – + +
CNQX – – +

**Fig. S5. Brain penetrant GABAB receptor antagonist induces spontaneous synaptic transmission on BrM2 cancer cells. A** and **B**. Example trace and quantification of sPSC frequency in BrM2 cells co-cultured with neurons following CGP 35348 and CGP + CNQX perfusion. Statistical significance was assessed using One-way ANOVA with Tukey's multiple comparisons (B).

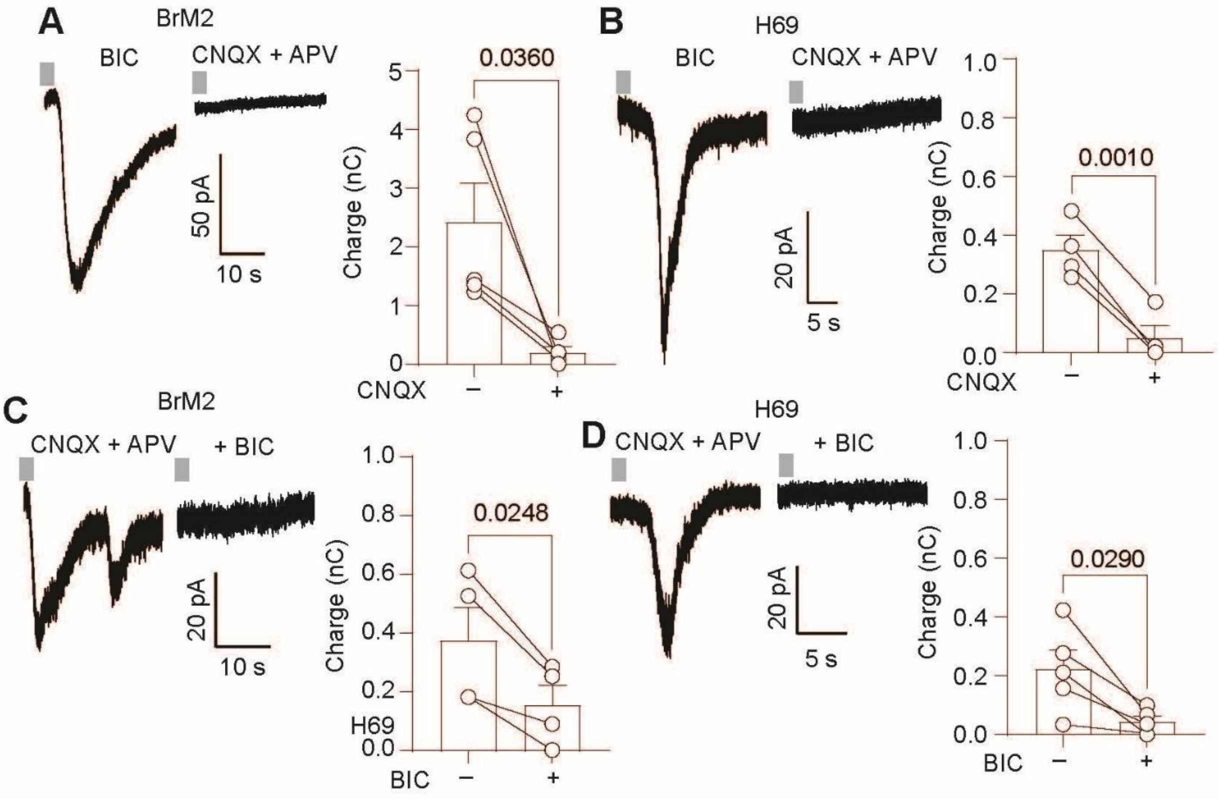
A
BrM2
BIC
CNQX + APV
50 pA
10 s
0.0360
Charge (nC)
CNQX
B
H69
BIC
CNQX + APV
20 pA
5 s
0.0010
Charge (nC)
CNQX
C
BrM2
CNQX + APV
+ BIC
20 pA
10 s
0.0248
Charge (nC)
H69
BIC
D
H69
CNQX + APV
+ BIC
20 pA
5 s
0.0290
Charge (nC)
BIC

**Fig. S6. H69 and BrM2 cells secrete glutamate and GABA. A.** Example trace and quantification of glutamate charge in neurons from BrM2 conditioned Tyrode perfusion. **B.** Example trace and quantification of glutamate charge in neurons from H69 conditioned Tyrode perfusion. **C.** Example trace and quantification of GABA charge in neurons from BrM2 conditioned Tyrode perfusion. **D.** Example trace and quantification of GABA charge in neurons from H69 conditioned Tyrode perfusion. Statistical significance was assessed using paired samples t-test (A, B, C, D).
